# Protective threshold of a potent neutralizing Zika virus monoclonal antibody in rhesus macaques

**DOI:** 10.1101/2024.08.16.608231

**Authors:** Joseph P. Nkolola, David Hope, Ruaron Guan, Alessandro Colarusso, Malika Aid, Robert H. Carnahan, James E. Crowe, Dan H. Barouch

**Author notes:** Corresponding author: Dan H. Barouch, M.D., Ph.D., Center for Virology and Vaccine Research, 330 Brookline Avenue, E/CLS-1043, Boston, MA 02115, USA. Author (Joseph Nkolola), (David Hope), (Ruaron Guan), (Alessandro Colarusso), (Malika Aid), (Robert H. Carnahan), (James E. Crowe Jr.).

## Abstract

Zika virus (ZIKV) is a mosquito-borne flavivirus that caused a global pandemic in 2016-2017 with continued ongoing transmission at low levels in several countries. In the absence of an approved ZIKV vaccine, neutralizing monoclonal antibodies (mAbs) provide an option for the prevention and treatment of ZIKV infection. Previous studies identified a potent neutralizing human mAb ZIKV-117 that reduced fetal infection and death in mice following ZIKV challenge. In this study, we report exquisite potency of ZIKV-117 in a titration study in rhesus macaques to protect against ZIKV challenge. We show complete protection at a dose of 0.016 mg/kg ZIKV-117, which resulted in median serum concentrations of 0.13 µg/mL. The high potency of this mAb supports its potential clinical development as a novel biotherapeutic intervention for ZIKV.

**Importance:** In this study, we report the potency of the ZIKV-specific neutralizing antibody ZIKV-117 against ZIKV challenge in a titration study rhesus macaques. This high potency supports the further development of this mAb for ZIKV.

## Introduction

Zika virus (ZIKV) is a member of the Flaviviridae family of positive-stranded RNA viruses and was first isolated in a rhesus macaque in Uganda in 1947 and in humans in 1952^1, 2^. Although primarily transmitted by the Aedes aegypti mosquito^3^, human-to-human transmission can occur via sexual^4^, vertical^5^ and blood transfusion^6^ routes. ZIKV outbreaks were reported in Micronesia in 2007^7^, Oceania in 2013–2014^8^, Brazil in 2015–2017^9^, and many countries in the Americas in 2016-2017^10^ and was declared a public health emergency of international concern by WHO^11^. While 80% of persons infected with ZIKV are asymptomatic or mildly symptomatic, during the 2015-2017 Brazilian epidemic a causal link was established between ZIKV infection and microcephaly and other congenital malformations in pregnant women^12^, as well as the neurologic condition of Guillian-Barré syndrome in adults^13^.

Although Zika cases have decreased significantly in frequency since their peak in 2017, the possibility of future outbreaks underscores the need for better preparedness, including the development of vaccines and biotherapeutics^14^. A previous study reported the identification of a potent neutralizing human mAb ZIKV-117, which was isolated from an otherwise healthy individual with history of symptomatic ZIKV infection^15^. ZIKV-117 neutralized ZIKV strains belonging to African, Asian and American lineages and mediated reduction of tissue pathology, placental and fetal infection, and mortality in murine models of experimental infection^15^. More recently, a nanostructured lipid carrier delivering an alphavirus replicon encoding ZIKV-117 showed robust protection both as a pre-exposure prophylaxis and post-exposure therapy in mice^16^. The mechanism of neutralization afforded by the ZIKV-117 mAb involves binding to domain II of the E protein on the ZIKV surface and cross-linking E glycoprotein dimers, resulting in prevention of the rearrangement of E proteins necessary for low-pH mediated fusion^17^. In the current study, we assessed the potency of ZIKV-117 in rhesus macaques to protect against ZIKV challenge.

## Materials and methods

### Study design

24 rhesus macaques were housed at Bioqual, Rockville, MD, and the study was conducted in compliance with all relevant local, state, and federal regulations and was approved by the Bioqual Animal Care and Use Committee (IACUC). Six groups of rhesus macaques (Macaca mulatta) were intravenously (I.V.) infused on day-1 with ZIKV-117 mAb at doses of 2.0, 0.4, 0.08, 0.016, 0.0032, or 0 mg/kg. To assess protective efficacy against ZIKV challenge, all groups were challenged subcutaneously (S.Q.) on day 0 with 10^6^ viral particles (VP) [10^3^ plaque-forming units (PFU)] of ZIKV strain ZIKV-BR isolated from northeast Brazil^18, 19^.

### Pharmacokinetics

Serum levels of the ZIKV-117 mAb were monitored using a previously described human IgG specific enzyme-linked immunosorbent assay (ELISA)^20^. In brief, ELISA plates were coated overnight at 4°C with 1 μg/mL of goat anti-human IgG (H+L) secondary antibody (monkey pre-adsorbed) (Novus Biologicals) and then blocked for 2 hours. Serum samples were assayed at 3-fold dilutions starting at a 1:3 dilution in Blocker Casein in PBS (Thermo Fisher Scientific) diluent. Samples were incubated for 1 hour at ambient temperature and then removed, and plates were washed. Wells then were incubated for 1 hour with horseradish peroxidase (HRP)-conjugated goat anti-human IgG (monkey pre-adsorbed) (Southern Biotech) at a 1:4,000 dilution. Wells were washed and then incubated with SureBlue Reserve TMB Microwell Peroxidase Substrate (Seracare) (100 μL/well) for 3 min followed by TMB Stop Solution (Seracare) to stop the reaction (100 μL/well). Microplates were read at 450 nm. The concentrations of the human mAbs were interpolated from the linear range of concurrently run purified human IgG (Sigma) standard curves using Prism software, version 11.0 (GraphPad).

### RT-PCR

Plasma viral loads after ZIKV-BR challenge were monitored for 3 weeks using a previously established reverse transcription-polymerase chain reaction (RT-PCR) assay^19^. In brief, the wildtype ZIKV BeH815744 Cap gene was used as a standard and was cloned into pcDNA3.1+, and the AmpliCap-Max T7 High Yield Message Maker Kit was used to transcribe RNA (Cellscript, WI, USA). RNA was purified using the RNA clean and concentrator kit (Zymo Research, CA, USA). Ten-fold dilutions of the RNA standard were reverse transcribed and included with each RT-PCR assay. Viral loads were calculated as virus particles (VP) per mL. Assay sensitivity was 50 copies/mL.

### Modeling

Estimation of a protective threshold for ZIKV-117 prophylaxis was performed by comparing ZIKV-117 mAb concentration in serum at the time of challenge with the time-weighted average values for the change of sgRNA viral load in serum samples from day 1 to 10 after viral challenge. A fitting curve was estimated using the locally weighted scatterplot smoothing (LOWESS) method as previously described^21^.

## Results and Discussion

24 rhesus macaques received an I.V. infusion of 2.0 (N=3), 0.4 (N=4), 0.08 (N=5), 0.016 (N=3), 0.0032 (N=3), or 0 (N=6) mg/kg ZIKV-117 on day -1. Serum ZIKV-117 levels were determined through day 10 by ELISA and showed the expected biphasic profile involving a rapid distribution phase followed by a slower elimination phase (Figure 1). Peak mAb levels were observed on day 0, one day following mAb administration. Animals that received 2.0, 0.4, 0.08, 0.016, 0.0032, and 0 mg/kg ZIKV-117 exhibited mean peak concentrations of 16.4, 4.5, 0.9, 0.13, 0.006, and 0 μg/mL respectively on day 0 (Figure 1). Circulating ZIKV-117 levels were observed over 10 days in all groups that received ZIKV-117, except for the 0.0032 mg/kg group for which we only detected borderline ZIKV-117 levels on day 0. No ZIKV-117 levels were detected in the sham control group.

**Figure 1.**
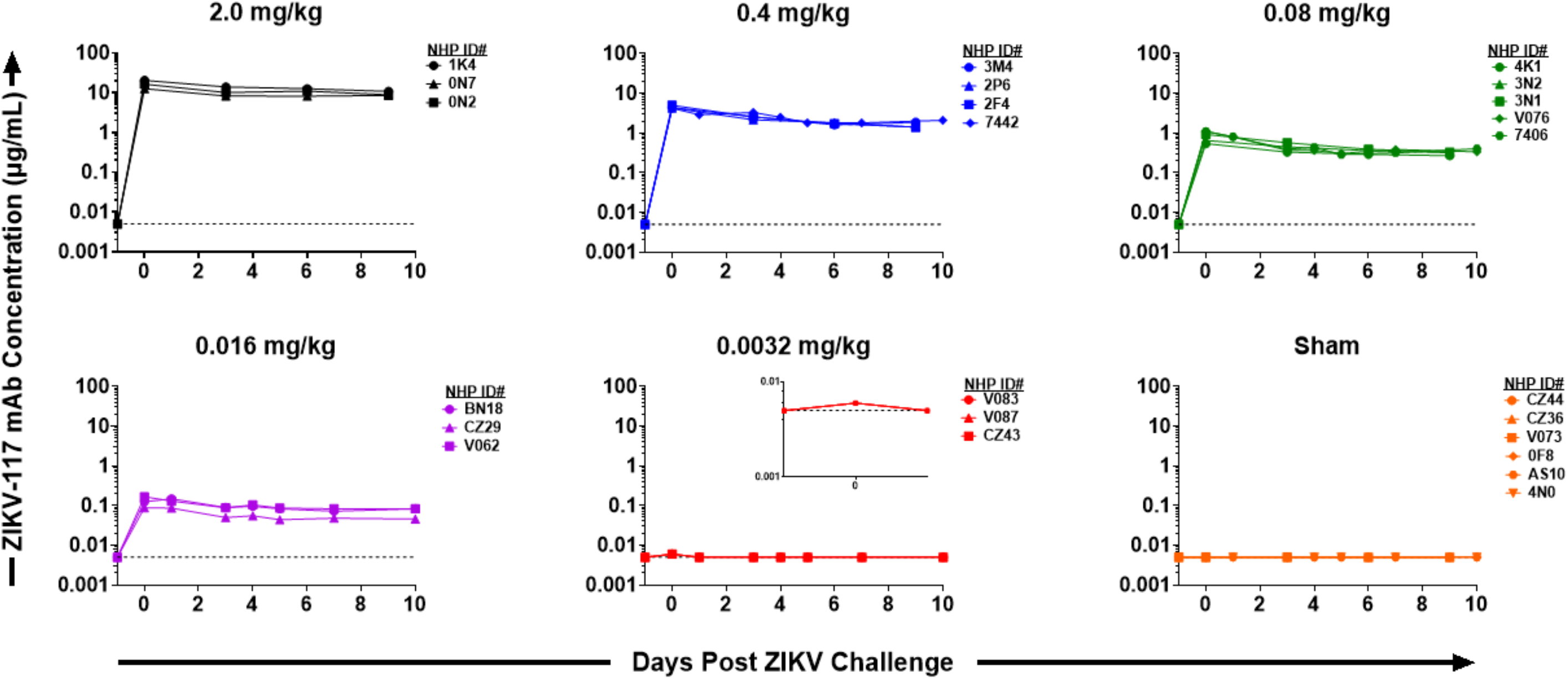
ZIKV-117 pharmacokinetics following infusion as measured by a human IgG-specific ELISA. Each line correspondences to a single animal. The horizontal dashed line represents the assay limit of detection (0.005 µg/mL).

All animals were challenged on day 0 with 10^6^ VP ZIKV-BR by the S.Q. route. The ZIKV-BR strain has been reported to recapitulate key clinical indications in wild-type SJL mice, including fetal microcephaly and intrauterine growth restriction^18^. Sham-inoculated animals demonstrated median peak viral loads of 4.7 log10 copies/mL (range 3.3 to 7.1 log10 copies/mL; n=6) on day 5-6 following challenge (Figure 2). In contrast, animals that received 2.0, 0.4, 0.08, and 0.016 mg/kg ZIKV-117 showed complete protection with no detectable viremia (<50 copies/mL) at all time points (Figures 2, 3A). Animals that received 0.0032 mg/kg ZIKV-117 showed median peak viral loads of 5.9 log10 copies/mL (range 2.2 to 6.5 log10 copies/mL; n=3) on day 6-7 following challenge, comparable to the sham control animals (Figures 2, 3A).

**Figure 2.**
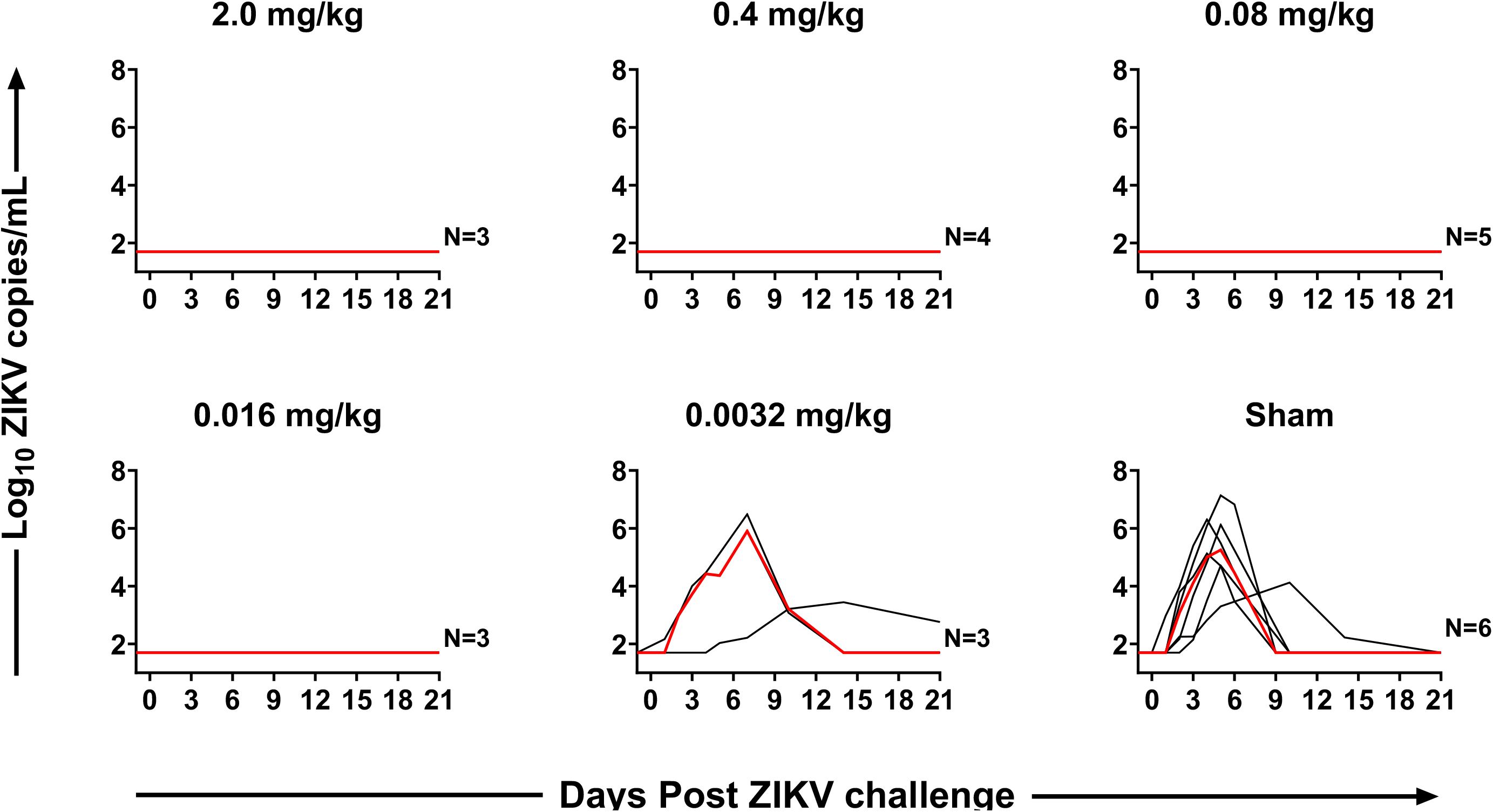
ZIKV-BR viral loads following challenge as determined by RT-PCR. Each black line corresponds to a single animal and the red line depicts the median for each group.

**Figure 3.**
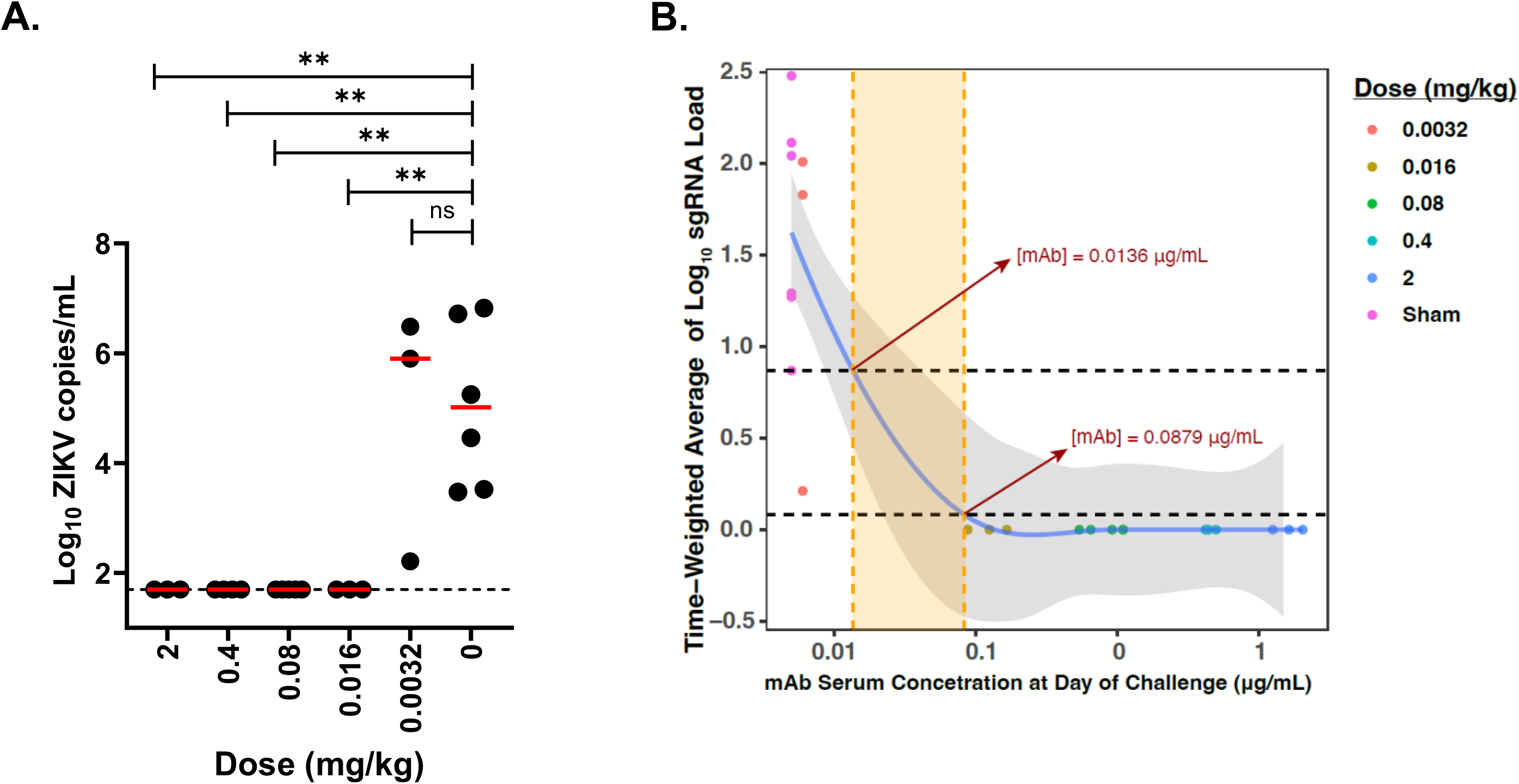
**(A)** Comparison of peak median ZIKV viral loads in each group compared with sham controls. The horizontal dashed line represents assay limit of detection (50 copies/mL). Adjusted Kruskal-Wallis tests are shown; **P<0.05. **(B)** Time-weighted average (TWA) values for the change of viral loads from day 1 to 10 after viral challenge (y-axis) compared with log serum ZIKV-117 concentrations on the day of challenge (x-axis). The fitting curve was estimated using the locally weighted scatterplot smoothing (LOWESS) method and is shown in purple, and gray shading indicates the 95% confidence interval. Horizontal black dotted lines indicate designated TWA thresholds for full (bottom line) and partial (top line) protection. Vertical dotted orange lines indicate maximal and minimal predicted cutoff for protective antibody concentration in serum.

We observed complete protection at a dose of 0.016 mg/kg ZIKV-117, which corresponded to median serum concentrations of 0.13 µg/mL (Figures 2, 3A). To model the threshold for protection more accurately, a fitting curve was estimated using the locally weighted scatterplot smoothing (LOWESS) method and suggested that the ZIKV-117 protective threshold was 0.014 – 0.088 µg/mL (Figure 3B). These data demonstrate the high potency of ZIKV-117 for protection against ZIKV-BR challenge in rhesus macaques (Figure 3B).

At least 89 countries and territories have had mosquito-borne transmission of ZIKV^22^. In this study, we show that ZIKV-117 is exquisitely potent for ZIKV protection in rhesus macaques. Doses of 0.016 mg/kg ZIKV-117 provided complete protection in this model, suggesting that 0.13 µg/mL is above the minimum protective threshold for this antibody. While a subset of potently neutralizing antibodies that target the conformational epitope of surface E glycoprotein dimers of both dengue virus and ZIKV have been identified^23^, ZIKV-117 remains the only ultrapotent ZIKA-specific neutralizing mAb to the best of our knowledge^24^. Taken together, our data suggests the potential of ZIKV-117 as a novel biotherapeutic for ZIKV.

## Acknowledgements

Authors would like to thank Camille Mazurek (BIDMC) for technical assistance.

## Author contributions

The study was conceptualized by R.H.C., J.E.C, and D.H.B; J.P.N., D.H. and R.G. performed the experiments; J.P.N., M.B. and A.C. performed data analyses; J.P.N and D.H.B. drafted the original manuscript. All co-authors contributed to reviewing and editing the final manuscript.

## Correspondence

Correspondence and requests for materials should be addressed to D.H.B.

## Funding

This study was supported by Defense Advanced Research Projects Agency (DARPA) grant HR0011-18-2-0001.

## Conflicts of Interest

J.E.C. has served as a consultant for Luna Labs USA, Merck Sharp & Dohme Corporation, Emergent Biosolutions, a former member of the Scientific Advisory Boards of Gigagen (Grifols), of Meissa Vaccines, and BTG International, is founder of IDBiologics and receives royalties from UpToDate. The laboratory of J.E.C. received unrelated sponsored research agreements from AstraZeneca, Takeda Vaccines, and IDBiologics during the conduct of the study. All other authors declare no competing interests.

